# A working memory intervention weakens the reconsolidation of a threat memory and its biases processing towards threat

**DOI:** 10.1101/2020.01.08.898643

**Authors:** Soledad Picco, Luz Bavassi, Rodrigo S. Fernández, María E. Pedreira

**Author notes:** These authors contributed equally to this work (Rodrigo S. Fernández, María E. Pedreira). Correspondence concerning this article should be addressed to Maria E. Pedreira, IFIByNE (CONICET), Ciudad Universitaria, (1428), Buenos Aires, Argentina.

## Abstract

**BACKGROUND:** Threat-conditioning (TC) memory plays a central role in anxiety disorders, but not in a simple way. This memory impacts on complex cognitive systems by modifying behavioral responses with a bias to fearful stimuli and overestimating potential threats. In this study we proposed a global approach analyzing the scope of disrupting TC memory reconsolidation in the implicit memory, the declarative contingency and the cognitive biases.

**METHODS:** Day 1: Subjects were trained on TC. Day 2: after Threat-memory reactivation, one group performed a high demanding working memory task (HWM) and the other a low demanding working memory task (LWM). The last group, only performed the HWM task. Day 3: TC memory was tested by an extinction session followed by reinstatement. Finally, all subjects completed tasks targeting stimuli representation, valuation and attentional bias towards threat.

**RESULTS:** Disrupting reconsolidation of TC memory with a HWM weakened the implicit memory retention and faded the representation and valuation towards threat but it had no effect on attentional bias. Moreover, we revealed that subjects’ performance during the working memory task was specifically associated in TC memory retention.

**CONCLUSIONS:** Our findings reveal a strong impairment of the threat-memory restabilization and associated emotional biases. This may result from the competition between defensive survival and central-executive control networks. Our results fits with Experimental Psychopathology approach, disentangling the relation between the implicit memory, cognitive, valenced systems and the possibility to weaken both the threatening memory and the systems associated with the maintenance of anxiety profile.

## INTRODUCTION

Anxiety disorders are the most frequent type of mental disorders being a relevant public health problem, which implies disability in the daily life and causes individual and societal cost (1,2). In this context, experimental models are necessary to understand deeply the nature of them. Considering that fear and anxiety are in the core of these disorders, it is possible to model such symptoms in animals and humans using conserved defense-survival responses (3). In the laboratory, the most useful paradigm is Threat-Conditioning (TC) (4). Thus, specific fear memories can be formed using the repeated pairing of an initially neutral cue (conditioned stimulus, CS) with a naturally aversive stimulus (unconditioned stimulus, US). As a result, an associative threat-memory is acquired. The later presentation of the CS will retrieve the US representation and elicit a conditioned threat response which predicts the occurrence of the US. Threat memory, plays a central role in anxiety disorders but not in a simple way (5,6). The CS predictive value for the negative outcome of the US, involves other components such as exacerbated negative representations associated with anticipated catastrophic consequences (7). The major concern, assuming this oversimplification, relies on the fact that, these defense-survival mechanisms are only one piece of the complex puzzle that contributes to human anxiety and neglects the complex cognitive-behavioral anxiety responses such as biased processing (8–10) towards fearful stimuli and overestimation potential threats(7,11,12).

In the last years, the immutable nature of consolidated memories has been challenged (13–15). After being consolidated, the presentation of a reminder (memory reactivation) of an acquired representation, opens a new round of re-stabilization of memory called reconsolidation (16,17). The reconsolidation process enables the update of consolidated memories. After its re-emergence in the field, a frequent proposal appears: the use of the reconsolidation process to modify maladaptive memories (18–21). Thus, different reports reveal that pharmacological interference of memory re-stabilization “erase” the implicit/emotional component of memory leaving the declarative component intact (22,23). This approach also showed to be efficient with a subclinical phobic sample and patients with post-traumatic stress disorders (24,25). A complementary approach, is the use of behavioral interference of memory reconsolidation. For example, the reactivation-extinction procedure uses extinction learning after memory reactivation (26). It was observed that, the negative value of the CS could be weakened (updated) with repeated presentations of the CS alone during the time window of the reconsolidation process. Thus, memory updating by the reconsolidation process reveals its effect on the implicit memory and maintains the declarative component unaffected.

A novel approach in the field suggests that memory reconsolidation depends on active working-memory processing of the reactivated memory (27). The circuits controlling working memory are limited and compete for resource allocation (28). Thus, if a working-memory intervention engages similar resources as those required for memory re-stabilization, the target memory is weakened as a result of the competition. For example, James and coworkers used the visuo-spatial game *Tetris* as an interference task during the reactivation of an intrusive film memory (27). The competition for visuo-spatial resources impaired the reconsolidation of visual features of intrusive memory. However, using a different task, where the target memory and the interference task, depended on the amygdala-based processing, Chalkia and coworkers(29) obtained negative results. That is, the execution of an emotional working memory task after threat-memory reactivation did not impaired memory reconsolidation. A possible explanation for these disparate findings can emerge from fMRI studies. With this technique, it was possible to describe large-scale functional networks, defined by a set of cortical and sub-cortical regions that co-activate together. For example, the salience network responds to threat and also to salient stimuli (30,31). Another described network is central-executive control network. Activation of this network, is observed during cognitively demanding tasks such as working memory maintenance (32). In a recent report it was shown that during acute threat the salience/defensive survival network and the central executive control network act as opposing systems (33).

Anxiety disorders engage physiological, emotional and cognitive changes (34). Consequently, focusing only on the analysis of acquired associative responses at physiological level (i.e. electrodermal activity or startle responses) disregards central features of this multifaceted phenomena, such as the cognitive and negative-valenced domains (35). To overcome these difficulties, we had developed a paradigm of TC, analyzing how this implicit memory could affect different valenced and cognitive systems in healthy participants (36). Subjects first underwent TC and then performed several tasks targeting biased cognitive processing towards threat (i.e. attentional bias) and the valuation systems (i.e. overestimation of threat and their consequences), which commonly underlie sub-optimal decision making in anxiety disorders (37,38). We found that participants rated as more aversive the reinforced stimuli(CS+) and generalized the fear to non-trained stimuli. Additionally, participants overestimated the probability of negative events associated with the CS+ and the reaction-times to CS+ showed an attentional bias towards threat. Overall, we found that TC in a healthy population could generate a similar biased processing towards threats such as those encountered in anxiety disorders.

Thus, we moved a step forward and analyzed the effect of reconsolidation interference of a threat-memory on the negative-valence and cognitive system. Here, we manipulated the demanding aspects of a working memory intervention, using a highly demanding working memory task (HWM, namely the Paced Auditory Serial Addition Task -PASAT: (39)) or a low demanding working memory task (LWM) as interference. We explored the interference effects not only on the implicit memory, but also on the negative valenced and cognitive systems. In this sense, we performed a TC protocol which included three groups. All subjects were trained on TC on Day 1, using male faces as CS’s. On Day 2, two groups were exposed to a cue-reminder and performed a HWM task or a low demanding working memory task (LWM, control intervention). The last group, only performed the HWM task. The TC memory was tested on Day 3 by the repeated presentation of the different CSs (extinction session) followed by the presentation of three unsignaled US’s (reinstatement). Finally, all subjects completed a set of different tasks targeting the negative-valenced and cognitive systems: stimuli representation (aversiveness) and threat-generalization, valuation (probability and cost) and attentional bias towards threat.

Our working hypothesis had two driven proposals. The first was that considering the competence between the salience and central executive networks for limited resources during memory reactivation, we expected that, the impairment of the TC memory re-stabilization would be associated with the demanding levels (high vs low) of the working memory intervention. The second, given the narrow relation between TC memory and the negative-valence and cognitive systems, its interaction might determine how reconsolidation interference may reduce acquired emotional and cognitive biases towards threat.

## METHODS AND MATERIALS

### Participants

A total of 67 undergraduate and graduate youths (37 females and 30 males) from the University of Buenos Aires (Argentina) mean age of 21,63 ± 2,69 were included in the study. Participants were randomly assigned to one of three groups. All participants gave written informed consent before the experiment, approved by Ethics Committee of the Review Board of the Sociedad Argentina de Investigación Clínica.

### Procedure

After arriving, subjects completed the subjective assessment questionnaires and were randomly assigned to a Reactivation-HWM (n=23), noReactivation-HWM (n=23) and Reactivation-LWM (n=21) group. On Day 1, participants underwent a TC. Day 2 consisted on a reactivation session of the TC followed by a working memory intervention (interference) or no-reactivation. On Day 3 an extinction was performed as a test of the retention of TC memory, followed by reinstatement. Finally, subjects performed different tasks to assess cognitive and negative-valenced domains. All tasks were programmed with Matlab (Mathworks Inc. Sherborn, MA, USA), Psychlab and the Psychtoolbox module.

### Subjective assessment

The State-Trait Anxiety Inventory (STAI-S and STAI-T, (35)) and the Beck Anxiety Inventory (BAI, (36)) questionnaires were used to asses participant’s trait-anxiety and the presence of physiological symptoms.

### Threat Conditioning

#### Acquisition

On Day 1, all participants underwent TC. Each face-stimulus (*see supplement*) appeared 8 times (24 trials total). One of the face-images (CS+) was followed by a tone-US (75% of reinforcement), while the other images (CS− and CSn) were not. All the CS’s were presented for 6s and the US appeared 1,5s before CS offset. The inter-trial interval varied between 8-12 s.

#### Reactivation

On Day 2, two groups (Reactivation-HMW and Reactivation-LMW) received a reactivation session, which consisted of an unreinforced presentation of the CS+, followed by an interruption and a black screen indicating the end of the experiment. After memory-reactivation, groups performed a working memory task (HMW or LMW). The noReactivation-HMW only completed the HMW without memory reactivation.

#### Extinction and Reinstatement

On day 3, all groups performed a retention test. Extinction consisted in 12 unreinforced presentations of each CS. Then, during reinstatement, participants were exposed to 3 consecutive unsignaled US, followed by the presentation of each CSs.

Skin Conductance Response (SCR) was used as measure of the implicit memory (TC) throughout the experiment (*see supplement*).

### US expectancy

Declarative memory of TC was assessed using an external keyboard (Yes/No buttons) in which subjects were required to answer the US expectancy during each CS presentation. On each trial, subjects were instructed to press “YES” button if they expected the CS to be followed by the US (tone) or “NO” button if they expected otherwise.

### Working Memory Intervention

#### HMW

The *Paced Auditory Serial Addition Test* (PASAT) (34) was used as the cognitive demanding task (HWM) for the Reactivation-HWM and noReactivation-HMW groups. In the **HMW** a pre-recorded series of numbers (1 to 9) were presented through headphones and subjects were asked to add the last two numbers. The task consisted in 61 trials (additions) and numbers were presented with a 2s interval. Only answers given within the interval were considered as correct For example, if the digits presented were ‘2’, ‘4’ and ‘8’, the correct answer would be first ‘6’ and then‘12’.

#### LMW

For Reactivation-LMW group, the task consisted in the same pre-recorded audio as the HMW, but subjects were asked to answer within a 2s interval if the digits were even or odd.

The demanding aspects of the working memory interventions were analyzed trough arousal and performance measures. Thus, arousal was assessed using SCL and it is reported as the mean (μS) tonic signal during the tasks. In addition, the proportion of correct responses was used as a measure of working memory performance.

### Cognitive and negative-valenced systems

#### Stimuli Representation and Generalization

Before TC on Day 1, and after its evaluation on Day 3, subjects rated the aversiveness of 12 different pictures of males faces using a 0-8 scale (5 angry, 4 neutrals and the ones used as CS+, CS− and CSn; (31)). Pictures were randomly presented on the screen until a response was given with the keyboard. For each group, the mean difference score between post-pre TC, was considered as the Stimuli representation score for CS+ and CS− and the mean difference (post-pre) of neutral and negative faces was considered as the Stimuli Generalization score.

#### Valuation (Probability and Cost)

On day 3, subjects read in the screen 24 positive and 24 negative scenarios, involving either the CS+ or the CS− along its picture (3,4). For each scenario and CS, subjects first responded how likely was the hypothetical event (in a 0 to 8 scale) and then how good/bad it would be for them, using the same 0 to 8 scale. For example, a negative event could be: *“How likely it would be you having problems at work with HIM?”* and then: *“How bad it would be for you having problems at work with HIM?”*. A positive situation could be: *“How likely it would be HE taking you to the airport?”*, and then: *“How good it would be HE taking to the airport?”*. A probability score and a cost score were calculated separately as the mean score difference between positive and negative scenarios for CS+ and CS−.

#### Attentional bias

On Day 3, after the extinction task (Figure 1), subjects performed a pictorial dot-probe task involving the CS’s (36). First, a fixation cross was presented in the center of the screen for 500 ms, followed by a simultaneous display of two CS (left/ right) for 500 ms. Then CSs pictures vanished and a small white probe appeared in the previous location (left/right) occupied by one of the CS for 1000 ms. Participants were asked to press a button as fast as they could, according to where they have detected the white probe (left/right) using an external keypad. Four types of stimuli compounds were used: *1)* CS+ vs CS−; *2)* CS+ vs CSn; *3)* CS− vs CSn and *4)* two neutral new faces (*filler* trials). We considered *Congruent trials* when the probe appeared on the CS+ location (stimuli compounds *1* and *2*) or CS− (stimulus compound *3*) and *Incongruent trials* the remaining possibilities (probe on CS− or CSn on compounds *1* and *2*; and CSn on compound *3*). Probe position (left/right) and face location was counterbalanced. Subjects performed a total of 160 trials in two blocks of 80 (40 trials *per* condition). Mean reaction times (RT) from the dot-probe task was estimated *per* condition taking the different between incongruent and congruent trials. Errors of commission, omission or responses less than 150 ms or greater than 2000 ms were not included in analysis.

**Figure 1.**
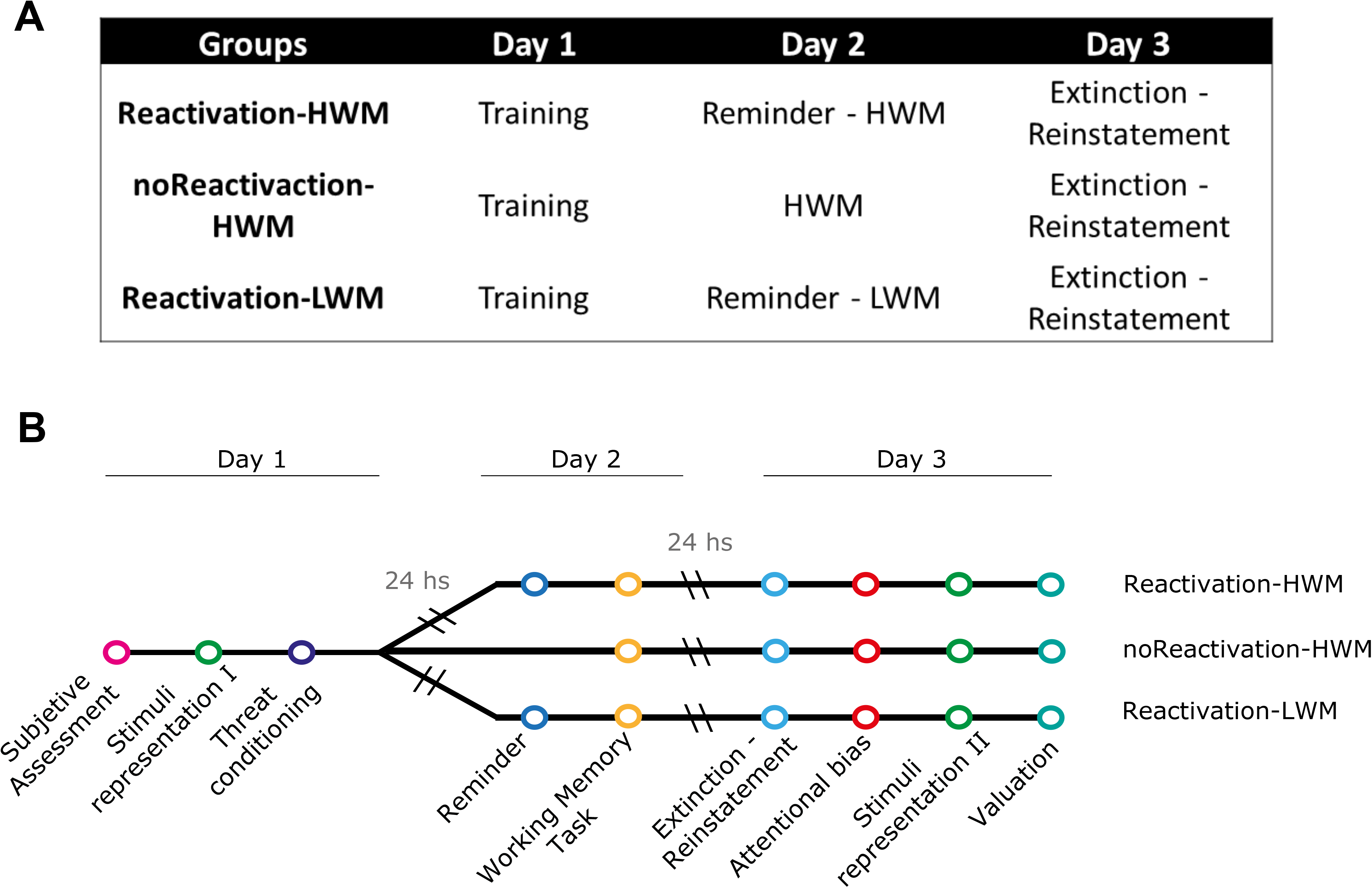
Schematic representation of the experiment. **(A)** Group description across the 3 Days. HMW stands for “Highly Demanding Working Memory Task” and LMW for “Low Demanding Working Memory Task”. **(B)** Experimental timeline for the 3 groups.

### Statistical Analysis

Subjective assessment and the proportion of correct responses during the working memory tasks (HWM and LWM) were analyzed using a One-Way ANOVA. SCR and proportion of YES responses for US expectancy during TC and extinction-reinstatement, were analyzed by means of a mixed ANOVA for repeated-measures with Group as the between-subjects factor and Stimulus (CS+, CS− and CSn) and Trial, as within-subject factors. When the interaction was significant, simple effects were performed. When sphericity was not accomplished, Greenhouse–Geisser correction was applied. Cognitive and negative-valenced systems were analyzed by separate Mixed-ANOVA’s (Group × Stimuli as factors), followed by post-hoc pairwise comparison using the Bonferroni correction for the main effects. Significant interactions were analyzed with simple effects and post-hoc Tukey comparisons.

## RESULTS

### Assessment

Participants did not differ in self-reported anxiety or working-memory capacity (*see supplement*, Table S1).

### Reconsolidation interference

Analysis of variance revealed a successful threat conditioning in all groups during acquisition on Day 1 (Figure 2; Mixed repeated-measures ANOVA, Stimulus: F_2.74_=15.994, p<0.001, ηp^2^=0.30; Stimulus × Trial Interaction: F_8.296_ =2.580, p<0.05, ηp^2^=0.06). There was a differential increase in SCR amplitudes for CS+ relative to CS− and CSn, from the first to the last trials of acquisition in all groups (last trial simple effects, CS+ vs CS_all_ p<0.001). As seen in Figure 2, a HWM after memory reactivation successfully interfered the re-stabilization of threat memory and prevented the return of fear. On Day 3, the group whose CS+ memory was reactivated and then performed the HWM task (Reactivation-HWM group) showed no differential SCR responding to none of the CS’s from the first trial of extinction (last trial of acquisition vs first extinction trial p<0.05, Group × Time interaction F_2.144_=2.063, p<0.05, simple effects, CS+ vs CS_all_ p>0.05). Moreover, the SCR presented no recovery after reinstatement (simple effects, CS+ vs CS_all_ p>0.05). In contrast, both the noReactivation-HWM and the Reactivation-LWM groups demonstrated threat-memory retention from Day 1 to Day 3 (simple effects, CS+ p>0.05) and differential responding to the CS+ in the first trial of extinction (simple effects, CS+ vs CS_all_ p<0.05). Moreover, the last trial of the extinction session showed no difference in SCR levels between CS+, CS− and CSn (p>0.05). Finally, both groups showed threat-memory recovery after reinstatement (simple effects, CS+ vs CS_all_ p<0.05). Consistent with previous results (23), reconsolidation interference of the implicit threat-memory did not affect the declarative component of this memory (*see supplement*, Figure S1, US expectancy).

**Figure 2.**
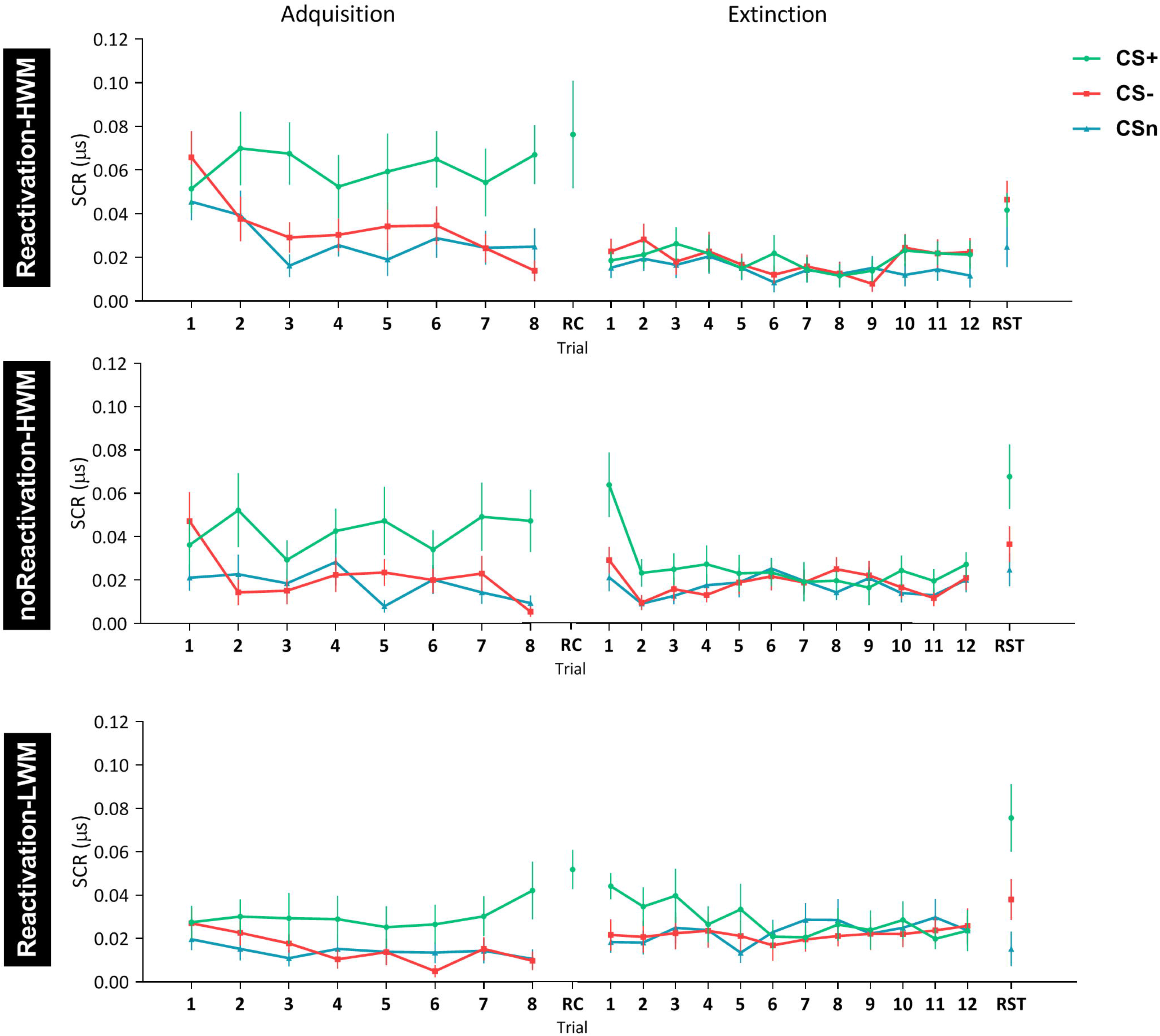
Reconsolidation interference of Threat Memory. **(A)** Reactivation-HWM group. **(B)** noReactivation-HWM. **(C)** Reactivation-LWM. Mean SCR (μS) ± SEM. Left panel: Threat conditioning acquisition on Day 1. Middle panel: Reactivation (RC) on Day 2. Right panel: Extinction training and reinstatement (RST) on Day 3. *P < 0.05.

### Working-Memory Intervention

Electrodermal activity (SCL) and subject’s performance, confirmed the cognitive-demanding nature of the HWM. On Day 2, groups that received the HWM (Reactivation-HWM and noReactivation-HWM) exhibited higher arousal (SCL levels) (Figure 3B, F_2_=4.154, p<0.001, ηp^2^=0.190) and had a worse performance during the task (F_2_=43.40, p<0.001, ηp^2^=0.583), relative to the group which performed the LWM (Reactivation-LWM). In order to disentangle the effect of the working memory intervention on the reactivated memory, we performed a correlation analysis including Reactivation-HMW and Reactivation-LMW (Figure 3C). As observed in Figure 3D, arousal and performance during working memory interventions were negatively correlated (r=−0.37, p<0.05), as high levels of SCL were associated with lower proportion of correct responses. Notably, subject’s performance predicted changes in threat memory retention between Day 3 and Day 1 (mean SCR difference between first extinction trial and last acquisition trial), (Figure 3E, r=0.41, p<0.001). Finally, we found no connection between arousal and changes in threat memory retention.

**Figure 3.**
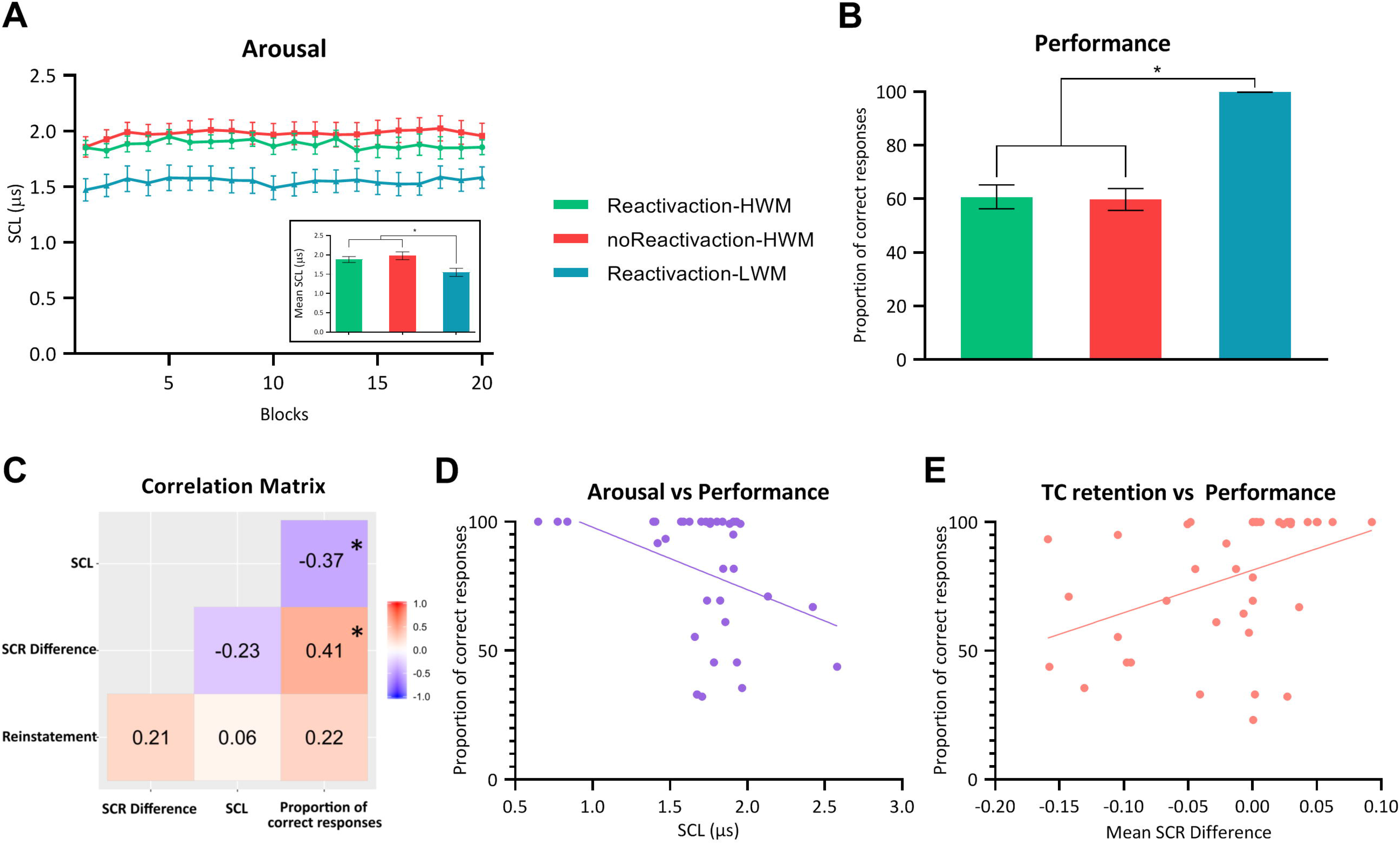
Subjects arousal and performance during the Working Memory Intervention (Day 2). **(A)** Mean SCL (μS) ± SEM across time for each group during the HWM or LWM. **(B)** Proportion of correct responses (± SEM) for each group during the HWM or LWM. **(C)** Correlation Matrix between the working memory interventions on Day 2 (SCL and Proportion of correct responses) and TC memory on Day 3 (Mean SCR difference and Reinstatement) **(D)** Arousal vs Performance correlation. **(E)** TC retention vs Performance correlation. *P <0.05.

### Cognitive and negative-valenced systems

#### Stimuli Representation

After threat-memory evaluation on Day 3, the Reactivation-HWM group, rated similarly the aversiveness of the CS+ and the CS− (Figure 4, Stimulus F_1.64_=12.074, p<0.001, ηp^2^=0.159, pairwise comparison p>0.05). In contrast, the noReactivation-HWM and the Reactivation-LWM groups, represented the CS+ significantly more aversive and unpleasant than the CS− (Figure 4, p<0.05 for both groups).

**Figure 4.**
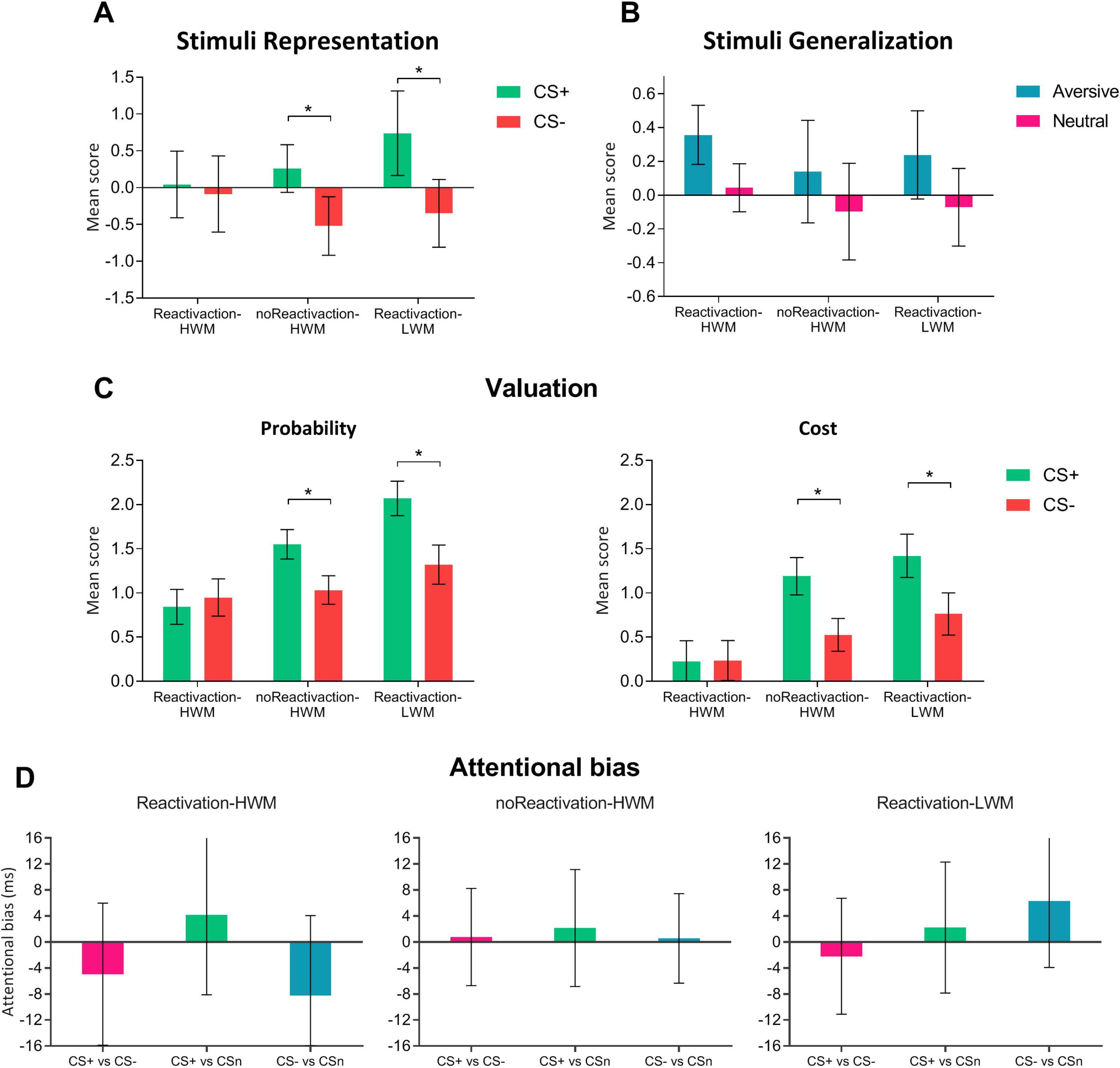
Cognitive and negative-valenced systems evaluation after threat conditioning testing (Day 3). **(A)** Stimuli Representation (aversiveness). Mean score (± SEM) difference between CS+ and CS−. **(B)** Stimuli Generalization. Mean score (± SEM) for non-trained negative and neutral pictures. **(C)** Valuation: Probability (left panel) and Cost (right panel). Mean score (± SEM) difference between positive and negative scenarios for CS+ and CS−. **(E)** Attentional bias (dot-probe). Difference in ms. between incongruent and congruent trials for 3 types of stimuli compounds (CS+ vs CS−, CS+ vs CSn and CS− vs CSn). *P <0.05.

#### Stimuli Generalization

Subjects rated the aversive faces higher than the neutral faces (Stimulus F_1.64_=4.009, p<0.05, ηp^2^=0.05), but we did not found difference between stimulus for each group (Figure 4, Stimulus × group F_2.64_=0.031, p>0.05, ηp^2^=0.001).

#### Valuation

A Mixed-ANOVA revealed that the Reactivation-HWM group, assigned a similar probability (Stimulus F_1.728_=7.156, p<0.01, ηp^2^=0.01) and cost (Stimulus F_1.729_=6.698, p<0.01, ηp^2^=0,009) to CS+ and CS− (pairwise comparison p>0.05) in either positive or negative scenarios (Figure 4). On the contrary, noReactivation-HWM and Reactivation-LWM groups, estimated higher probability to the CS+ relative to the CS− (p<0.05 for both groups) and its associated cost (p<0.05 for both groups).

#### Attentional bias

We found no evidence of attentional bias toward threat in the dot-probe task. A Mixed-ANOVA revealed a similar performance in all groups and no difference between them (Figure 4, Stimulus F_2.108_=1.433, p>0.05, ηp^2^=0.02; Stimulus × Group F_4.108_=0.89, p>0.05, ηp^2^=0.003). Moreover, RT did not differ from zero in any condition (*see supplement*, Table S2).

## DISCUSSION

Threat-conditioning (TC) is generally an adaptive and self-preserving form of learning which generates an implicit memory. However, it may become a source of pathologies when the reactivity to threat-cues persists in the absence of the reinforcement (7). Initially, this type of learning has been associated with development of anxiety disorders (6,40). Different pieces have been added to the puzzle, to understand this type of disorders such as incubation of fear, failure to inhibit the fear response to safety cues and stimulus generalization (37). Considering this background, in the present report, we demonstrated how the impairment of TC memory re-stabilization by a HMW, affected the implicit memory but not the declarative memory (Figure S1). In addition, we revealed that not any working memory task interfere memory reconsolidation because the interference effect depends on the level of cognitive demand (Figure 3) (28). Finally, we showed how the interference of memory reconsolidation reduced biased processing towards threat (i.e. stimuli representation and valuation). Figure 4).

Previously, we demonstrated that attentional bias towards threat could be acquired and maintained at least 48 h after TC, using the dot-probe task, which is commonly used to assess biased processing (41). However, in the present design, we included before the dot-probe task, an extinction training and memory reinstatement. These manipulations after memory consolidation, may be responsible of the lack of an attentional bias as reported in Fernández and colleagues (36). In this sense, other authors showed that the number of extinction and reinstatement trials should be similar in order to recover the attentional bias. This premise was not fulfilled in the present experimental design, as we used 12 extinction trials and only 3 reinstatement trials.

Few reports used cognitive-interventions to engage and overload working-memory in reconsolidation studies. First, James and co-workers used an episodic memory based on “traumatic film”. After memory-reactivation, subjects played a computer game (*Tetris*). They found a diminution in the number of memory intrusions of the “trauma film”, compared with the control group. The target memory and the interfering task, required visuo-spatial working memory, thus the competition for similar resources may explain the weakened of the visual aspects of the intrusive memory. Using a similar target memory, Gotthard and Gura (42) used a video to create a positive declarative memory. After its reactivation, participants performed a visuospatial interference task (word search). This interference was effective in reducing free recall of the video details at testing. Finally, Chalkia and coworkers (28), found that an emotional working memory task with visuo-spatial processing, was unable to impair the re-stabilization of a threat memory. They proposed, that the negative results may depend on an inappropriate or low demanding working memory task. Here, regardless of the different nature of the threat-memory and HWM, we found a strong impairment on memory re-stabilization. The whole understanding of these finding, may arise from taking into consideration the different large-scale functional networks. The salience/defensive network shows similarities with the defensive survival circuits (3). Moreover, another large-scale network, is the central-executive control network, associated with cognitively demanding tasks such as working-memory maintenance and the reduction of subjective feelings of fear and anxiety (32,43). Different reports, suggested that the salience/defensive survival-network and the central executive control network act as opposing systems (44). When participants perform a working-memory task during a threatening situation, activation in the central-executive control network is reduced compared to a nonthreatening context. In line with these results, other report showed that the connectivity within the salience/defensive survival network increase during acute threat and decrease in the central-executive control network. Behaviorally, it was demonstrated that during TC acquisition is impaired, when simultaneously subjects perform a demanding working memory task(45).

This context supports the results obtained in this report. A HMW as the PASAT, may compete for neural resources impairing the threat-memory reconsolidation. We demonstrated that, the cognitive effort but not arousal levels during the HWM (Figure 3), may be related to the postulated effect of the salience/defensive survival-network and the central executive control-network on memory re-stabilization. Finally, the characteristics of our experimental design with healthy participants are in line with two main translational proposals in the field. First the use working-memory tasks to interfere emotional memories re-stabilization. Different interventions have been candidates such as the Eye Movement Desensitization and Reprocessing, *Tetris game* and the cognitive reappraisal technique, showed positive effects on emotional memory weakening (27,46). However, the ideal candidate should consent to increase or decrease the cognitive demand. In this report, we showed that the cognitive effort of the working-memory task, could be monitored by *online* measures such as sympathetic activity. In addition, this effortful aspect of the working-memory task could be adjusted according to the protocol requirements such as the time interval between stimuli. To conclude, considering the unique complexities of anxiety disorders, it is necessary to add to animal models of anxiety disorders and clinical research with patients, the valuable resource of studies in healthy subjects. It offers a new inside named Experimental Psychopathology approach (47). This approach relies on a continuum from normal to pathological anxiety and considers the underlying mechanisms. Thus, the results obtained in this study disentangle the relation between the implicit memory, cognitive, valenced systems and the possibility to weaken both the threatening memory and the systems associated with the maintenance of anxiety profile. Future research may reveal if these effects also may be obtained in subjects with different anxiety profiles.

## ACKNOWLEDGEMENTS AND DISCLOSURES

We thank Angel Vidal for technical assistance. This work was supported by Agencia Nacional de Promoción Científica y Tecnológica (http://www.agencia.mincyt.gob.ar/) PICT 2013-0412 and PICT 2016-0243 (MEP). The funders had no role in study design, data collection and analysis, decision to publish, or preparation of the manuscript. The authors declare no conflict of interest.

