## Supplement for "A working memory intervention weakens the reconsolidation of a threat memory and its biases processing towards threat"

### METHODS AND MATERIALS.

#### Stimuli.

Three different fear relevant male faces pictures, taken from the Karolinska Directed Emotional Faces database served as the Conditioned Stimuli (CSs). They were presented in the center of a black screen (slides of 9,5 cm × 7 cm), two of them (CS+ and CS-) expressed anger and the other one (CSn) was neutral (no emotion). An auditory stimulus (tone) with a duration of 1,5 s delivered through stereo headphones served as the unconditioned stimulus (US). The tone was generated by a TG/WN Tone-Noise Generator (*Psychlab*), digitally controlled with a mean of 98 db±4 db. The US was adjusted for each subject to be "unpleasant but not painful" (100 db was the maximum allowed for any subject). Before TC, 3 habituation trials were performed. Assignment of the aversive pictures (CS+ and CS-) as the reinforced stimulus was counterbalanced across subjects.

**Electrophysiological measures.**

*Skin Conductance Response (SCR) and Skin Conductance Level (SCL).* Electrodermal activity was measured using an input device (*Psychlab* Precision Contact Instruments) with a sine excitation voltage (±0,5 V) of 50 Hz derived from the main frequency. The device was connected to two Ag/AgCl electrodes of 20 mm × 16 mm located in the intermediate phalanges of the non-dominant hand. The raw SCR scores were square-root transformed to normalize distributions. A minimum response criterion of 0,002 micro Siemens (μS) was used and all the other responses were scored as zero. Only subjects who showed differential fear responding (CS+ SCR amplitude > CS- and CSn) were considered for analysis. SCL was used to measure arousal and the demanding cognitive aspects of the HMW and LMW and it is reported as the mean (μS) tonic signal during the tasks.

### RESULTS.

**US expectancy (Declarative memory):**

At the end of acquisition, subjects showed a higher percentage of YES responses associated to the CS+ presentation for all groups, related to the learned contingency between CS+ and US during TC. Proportion of YES responses for CS+ differ significantly from CS- and CSn in acquisition (trials 1-8) (Repeated-measures ANOVA, Stimulus × Time

Interaction: $F_{2.90}$ =320.951, p<0.001, ηp2 =0.87; simple effects CS+ vs $CS_{all}$ p<0.001). Moreover, US expectancy related to CS+ was maintained in the first trial of extinction, on Day 3 as well. During extinction session the percentage of YES responses associated with the presentation of the CS+ significantly decreased (proportion of YES responses extinction session trials 1-12, simple effects p<0.001). Finally, in reinstatement session, percentage of YES responses increased in all groups.

| Groups | Acquisition<br>Trial 8 | Extinction<br>Trial 1 | RST |
|---|---|---|---|
| **Reactivation-HWM** | 100% | 62.50% | 62.50% |
| **noReactivaction-HWM** | 100% | 66.66% | 56.00% |
| **Reactivation-LWM** | 94.73% | 83.33% | 54.00% |

**Percentage of YES response to CS+ across groups.**

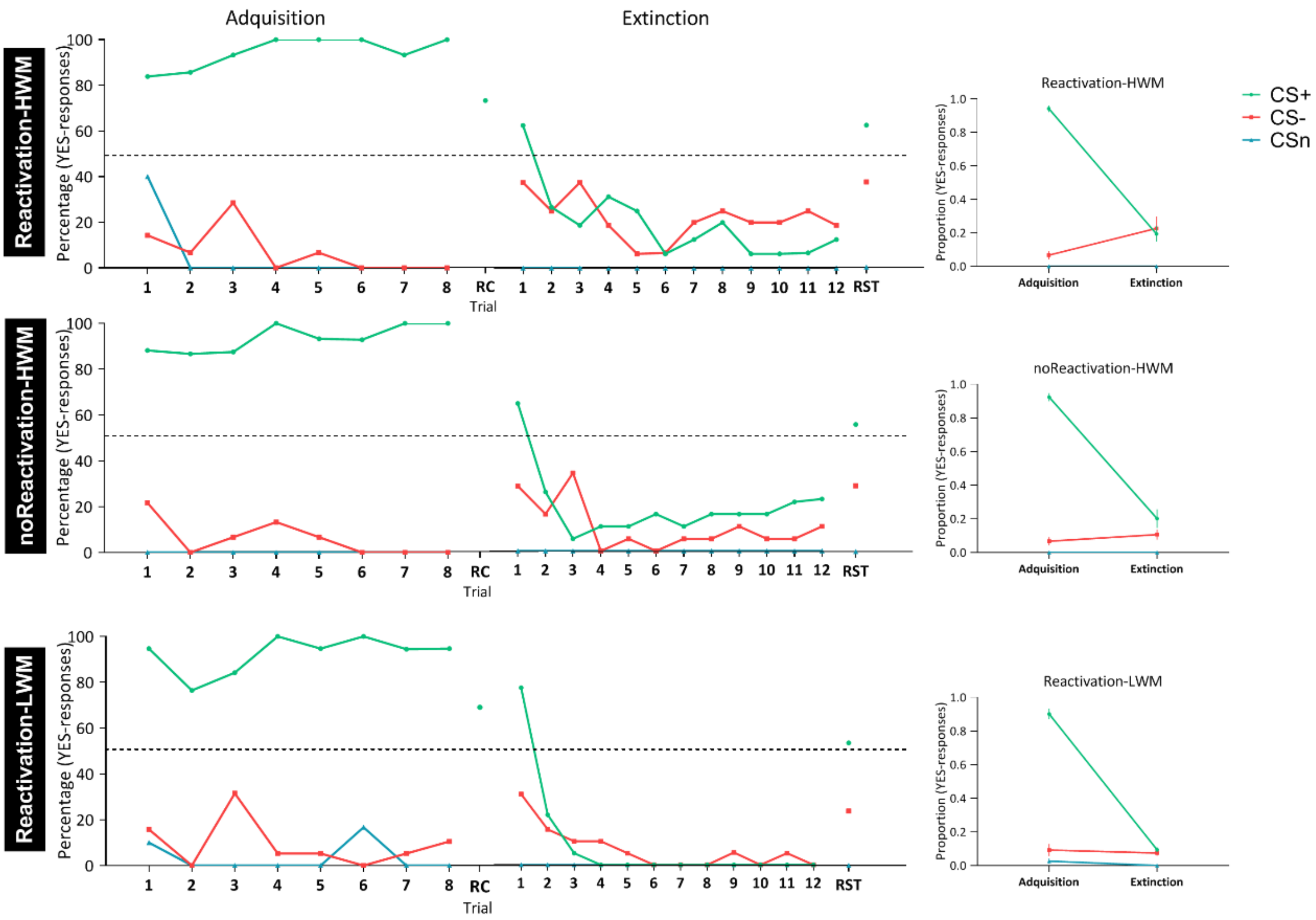


**Figure S1. US expectancy during TC. (A)** Reactivation-HWM group**. (B)** noReactivation-HWM. **(C)** Reactivation-LWM. Left panel: Percentage of YES responses during Threat conditioning acquisition on Day 1. Middle panel: Percentage of YES responses during Extinction training and reinstatement on Day 3 (RST). Right panel: Proportion of YES responses for acquisition (trial 1-8) and extinction (trial 1-12).

**Table S1. Subjective Assessment (Day 1).**

**Table S1. Subjective Assessment (Day 1).** Mean values ± SEM of the State-Trait Anxiety Inventory (STAI-S and STAI-T), Beck Anxiety Inventory (BAI) and Working Memory Capacity (PASAT) for each group.

| Groups | Reactivation-HWM | noReactivaction-HWM | Reactivation-LWM | Anova Results |
|---|---|---|---|---|
| **STAI-T** | 32.00 ± 1.40 | 31.61 ± 1.46 | 33.76 ± 1.77 | $F_2$=0.536, p>0.05 |
| **BAI** | 9.65 ± 0.93 | 9.17 ± 0.62 | 10.43 ± 0.93 | $F_2$=0.561, p>0.05 |
| **STAI-S** | 34.57 ± 1.26 | 32.43 ± 0.93 | 34.14 ± 1.75 | $F_2$=0.737, p>0.05 |
| **WM Capacity** | 38.61 ± 2.84 | 38.18 ± 2.62 | 60.90 ± 0.07 | $F_2$=0.197, p>0.05 |

**Table S2. Attentional bias. Statistical analysis.**

| Groups | CS+ vs CS- | CS+ vs CSn | CS- vs CSn |
|---|---|---|---|
| **Reactivation-HWM** | $t_{19}$=-1.633, p=0.119 | $t_{19}$=-1.740, p=0.098 | $t_{19}$=-0.076, p=0.940 |
| **noReactivaction-HWM** | $t_{19}$=-0.284, p=0.780 | $t_{19}$=-0.098, p=0.923 | $t_{19}$=0.846, p=0.408 |
| **Reactivation-LWM** | $t_{16}$=-0.049, p=0.961 | $t_{16}$=-0.712, p=0.486 | $t_{16}$=-1.285, p=0.217 |

**Table S2. Attentional bias. Statistical analysis.** T-test comparison for each stimuli compound (CS+ vs CS-, CS+ vs CSn and CS- vs CSn) against zero.
